# RetSynth: Solving all optimal retrosynthesis solutions using dynamically constrained integer linear programming

**DOI:** 10.1101/223271

**Authors:** Leanne S. Whitmore, Ali Pinar, Anthe George, Corey M. Hudson

## Abstract

**Motivation:** Naive determination of all the optimal pathways to production of a target chemical on an arbitrarily defined chassis organism is computationally intractable. Methods like linear integer programming can provide a singular solution to this problem, but fail to provide all optimal pathways.

**Results:** Here we present RetSynth, an algorithm for determining all optimal biological retrosynthesis solutions, given a starting biological chassis and target chemical. By dynamically scaling constraints, additional pathway search scales relative to the number of fully independent branches in the optimal pathways, and not relative to the number of reactions in the database or size of the metabolic network. This feature allows all optimal pathways to be determined for a very large number of chemicals and for a large corpus of potential chassis organisms.

**Availability:** This algorithm is distributed as part of the RetSynth software package, under a BSD 2-clause license at https://www.github.com/sandialabs/RetSynth/

## 1 Introduction

Flux Balance Analysis (FBA) is a tool widely used in predicting metabolic behavior at genome scale [1]. A common application of FBA is the determination of pathways for synthetic genetic manipulation [2]. This may involve the addition of enzymes, usually by means of plasmids, or gene silencing. The result of this technique is the dynamic manipulation of an organism’s metabolic output [3].

FBA requires, at a minimum, a stoichiometric matrix (*S*). This matrix is ideally complete with regard to the available reactions and compounds for a given organism. The reactions, are conventionally tied to a set of explicit enzymes and transporters, but additional enzymes and transporters may be provisionally assigned as needed to provide a complete solution. Secondarily, an objective function (*Z*) is solved for the metabolism of interest. This may involve minimization of input, maximization of output, or other constraints [4]. Frequently biomass is an important characteristic of the objective function. This subfunction, is a mathematical device used to represent the metabolic requirements and products of the cell division or reproduction process [5].

A well understood problem in FBA, is that the results produce a single *υ* solution to *Sυ* = *b*, where many may exist. One way of circumventing this problem is flux variability analysis (FVA), which gives alternate optimal solutions [6]. These solutions are presented as ranges over the flux vector and no single optimal solution is presented in FVA. This method characterizes the range of available optimal fluxes. A similar problem is how to characterize alternative optimal pathways using Mixed Integer Linear Programming (MILP). Lee [7] and Phalakornkule [8] did early work on alternative optima in linear programming models of metabolism. The work that we will present draws inspiration from these models. However, our technique characterizes not solely the metabolic control schemes, but also the available strategies for optimal metabolic modification.

An interesting problem in synthetic biology, is determining the minimal number of gene additions that would be required to get an industrial organism to optimally produce a compound of interest. In the case where the organism natively produces the compound, non-optimally, this results in a classic FBA problem. A typical approach then, is to solve the problem twice, maximizing biomass in one solution, then maximizing biomass and the compound of interest in the second solution. Genes encoding enzymes controlling reactions with reduced flux in the second solution can be silenced. Assuming the biomass function is sufficiently complete, this will result in shunting precursor metabolites to reactions producing the compound of interest and limiting subsequent reduction of the compound, after it has been produced. Several caveats must be accepted here, including that the compound of interest does not create a toxic environment and that the stoichiometric matrix and biomass equations are sufficiently complete, so that necessary biomass precursors are not consumed by the reactions producing the compound of interest.

In cases where the organism does not natively synthesize the compound of interest (hereafter *x*), a less obvious optimization problem is needed to determine the optimal strategy for producing *x*. Naively, it is possible to manually add a single non-native reaction to the stoichiometric matrix. Here the non-native reaction is the assumed stoichiometry of a known enzymatically-controlled chemical reaction. If there are *k* reactions that produce *x*, the first step is to try each of the *k* reactions to see if biomass and *x* can be produced. If we want to know all reactions that could be added, we need to try all *k* reactions. If there is a single step solution, it solves in FBA(*k*) time. If there is no single step solution, the problem explodes exponentially, both in the difficulty and complexity. A two-step solution requires not just *k* reactions, but all reactions that produce the precursors to the *k* reactions. If the average number of reactions producing any given compound is 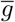 then the number of solutions that must be tested for a *y* step solution in the worst case is 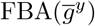.

In the description of the algorithm below, which alleviates the worst-case solution 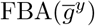, we show how given a database of potential reactions and a genome scale metabolic model, we can determine in a roughly linear number of FBA steps, all optimal genetic additions required to optimally produce a given compound of interest. This is done by dynamic and recursive manipulation of the constraint space, followed by integer programming. Setting a maximum number of reaction additions (pathway length constraints), the solvability of the problem can be efficiently determined. This technique requires only a genome scale metabolic model, a database of potential reactions and a compound of interest.

## 2 Algorithm

The algorithm below shows how to add reactions to a metabolic model, in order to determine all optimal additions. In general, optimal solutions are defined by the minimum number of reaction additions to solve a given problem. In cases where experimental flux can be determined, or enzyme productivity can be calculated, multiple optimal solutions can be ranked, based on output criteria.

The metabolic system is defined as:

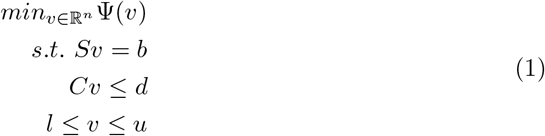

where 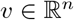 defines the rate of each reaction (i.e., flux), Ψ is convex and maps the space of potential fluxes to a single optimal solution, 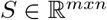 is the stoichiometric matrix *mxn*, with *m* molecules and *n* reactions and *b* is a vector of exchanges, such that *Sυ* = *b* and *b* is conventionally defined as 0, *l* and *u* are the lower and upper bounds of any given reaction, respectively, C is a matrix representing a system of equations resulting in a linear inequality w.r.t vector *d*.

The gene addition problem is defined as:

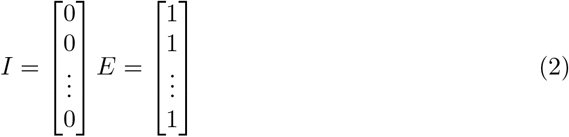

where *I* is a vector of constraints in *C* for natively produced chemical reactions in the organism of interest, and *E* is a vector of constraints in *C*^*^ for all possible chemical reactions. The 0-1 MILP problem is defined as:

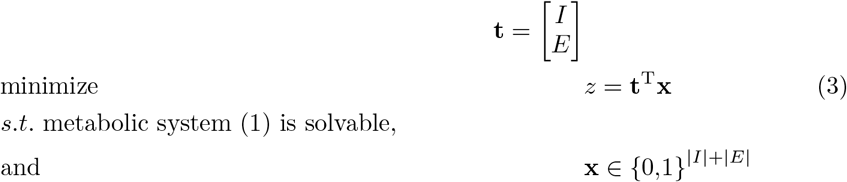

where *z* corresponds to the number of external reactions to be added to the system. This is solved by extracting rows from *C*^*^ where *x* = 1, appending them to *C* and solving (1).

The idea behind our algorithm is to add a penalty function to variables that are already identified as part of a solution to force the algorithm to seek alternative optimal solutions. We compute the penalty such that any optimal solution to the modified problem remains an optimal solution to the original problem. That is *t^T^x* < *β*^*^(1 + 1/(2*β*^*^) < *β*^*^ + 1. This implies that any enzyme added to 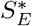 is part of an optimal solution.

We also need to show that the list is complete and any enzyme that is part of an optimal solution is in 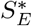. Assume the contrary, and let the *j*^th^ enzyme be part of an optimal solution but not included in 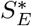. Then we have *t_j_* = 1. The *t* values for the other *β*^*^ – 1 enzymes that are part of the optimal solution can be at most 1 + 1/(2*β*^*^). All together the optimal solution value to the modified problem will be *β*^*^ + 1/2 – 1/(2*β*^*^). However, the algorithm terminates only after the optimal solution to the modified problem reaches *β*^*^(1 + 1/(2*β*^*^)), which is higher than the solution that includes the *j*^th^ enzyme. This leads to a contradiction and proves that our algorithm includes all enzymes that are part of an optimal solution.

### 2.1 Sub-optimal solutions

In addition to finding all optimal solutions, we extend this algorithm to find solutions that may involve more than the optimal number of added reactions. RetSynth is able to find solutions up to *β*^*^ + *k*, where *k* is a parameter set by the user. The modification of the algorithm involves setting *t* values to 1 + 1/(*β*^*^(*k* + 2)) when *t^T^x_i_* > 0 and repeating the loop until *t^T^x* ≥ (*β*^*^ + *k*)(1 + 1/(*β*^*^(*k* + 2))). After each increase in 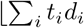, then 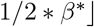, 1/(2 * *β*^*^ + 1) is added to *t_i_* where 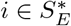. This allows a solution space that includes not only the optimal solutions, but all solutions < *β*^*^ + *k*. The correctness of the algorithm can be proven the same way.

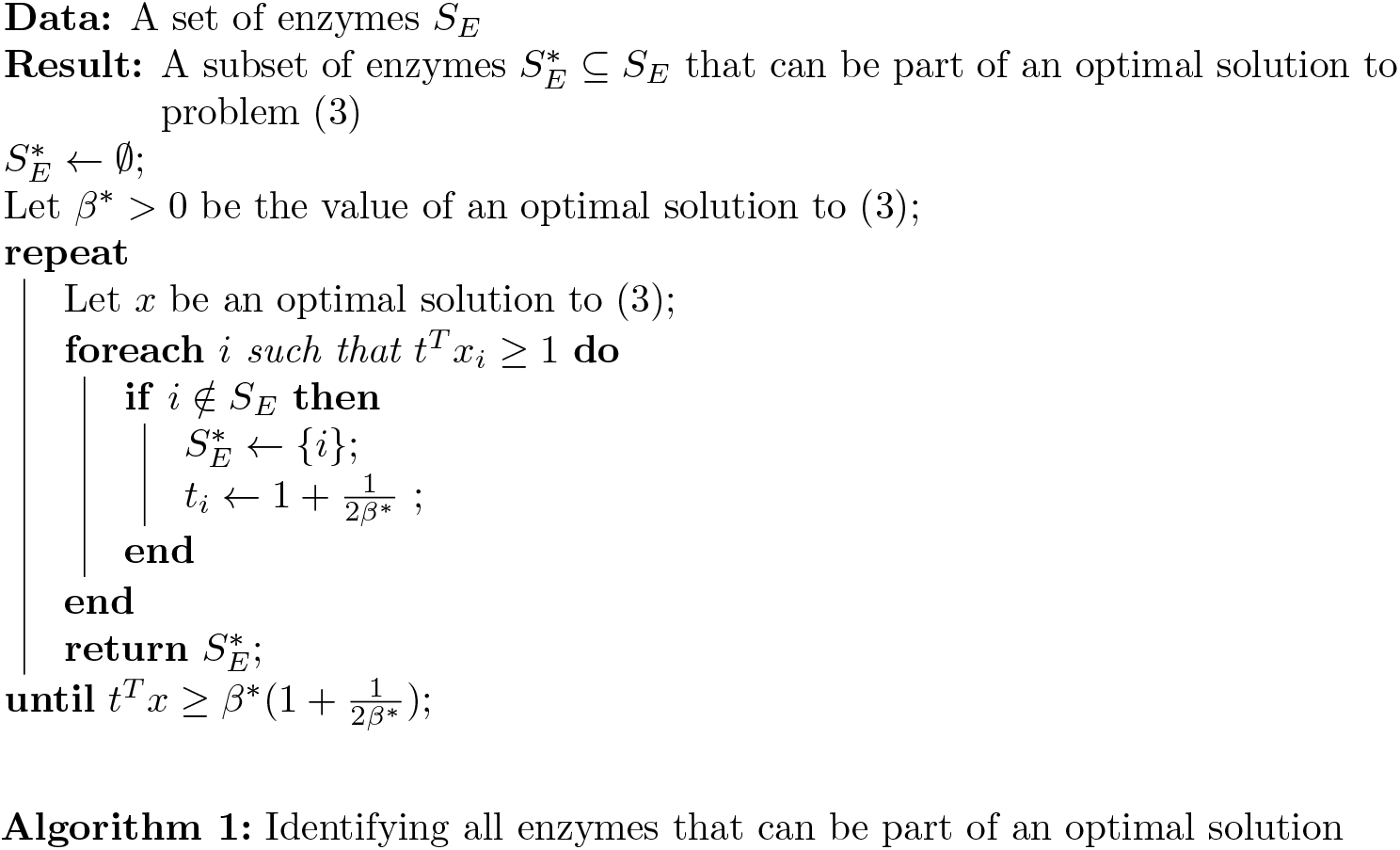

### 2.2 Enumerating and backtracking all solutions

The new set 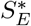 is typically much smaller than 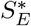, and drastically reduces the search space for enumerating all optimal solutions. Next, we will describe how this can be done.

Define a directed graph *G* = (*V, E*) with two types of nodes: *V* = *V_c_* ∪ *V_p_* and 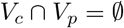. The process nodes,*V_p_*, represent the enzymes selected in the previous section, whereas the compound nodes, *V_c_*, represent all compounds that are inputs to the processes. The directed edges will represent the input/output relationships between compounds and processes. That is we will add an edge from a compound node to a process node, if this process requires the compound as an input. Symmetrically, we will add an edge from a process node to a compound node, if the process produces this compound. For completeness, we will add a super-process node for all processes that are already active, along with edges to all compounds these processes can produce.

Conjecture: *Given a compound of interest and a dependency graph G, a connected subgraph which includes the node for the compound of interest and at least one predecessor node for each compound node describes a feasible solution to our problem. Symmetrically, any feasible solution is a subgraph that satisfies these conditions. Subsequently, such a subgraph with minimum number of process nodes (not counting the super-process node) will define an optimal solution*.

Enumerating these results turns into a path enumeration problem, where we enumerate minimal length paths from the super-process node to the node for the compound of interest.

## 3 Software Description

This software has been distributed with a permissive open source license (BSD 2-clause), is written in Python and available for use on GitHub (http://www.github.com/sandialabs/RetSynth/).

### 3.1 Comparison with other software

A variety of algorithms exist for solving the retrosynthesis problem. These include BNICE [9], GEM-Path [10], DESHARKY [11], ReBIT [12], RetroPath [13], PathPred [15], RouteSearch [14] and SimPheny (www.genomatica.com). The difference between RetSynth and the other available algorithms, is that RetPath provides all available optimal and *β*^*^ + *k* solutions. The efficiency of the algorithm (scales by the number of independent pathways) allows this to be run for any chassis with a fully characterized metabolic network and for any target chemical.

## Acknowledgements

Sandia National Laboratories is a multi-mission laboratory managed and operated by National Technology and Engineering Solutions of Sandia, LLC, a wholly owned subsidiary of Honeywell International, Inc., for the U.S. Department of Energy’s National Nuclear Security Administration under contract DE-NA0003525.

## Funding

This research was conducted as part of the Co-Optimization of Fuels & Engines (Co-Optima) project sponsored by the U.S. Department of Energy (DOE) Office of Energy Efficiency and Renewable Energy (EERE), Bioenergy Technologies and Vehicle Technologies Offices. Co-Optima is a collaborative project of multiple national laboratories initiated to simultaneously accelerate the introduction of affordable, scalable, and sustainable biofuels and high-efficiency, low-emission vehicle engines.

